# Evolution of embryo implantation was enabled by the origin of decidual cells in eutherian mammals

**DOI:** 10.1101/429571

**Authors:** Arun R. Chavan, Oliver W. Griffith, Daniel Stadtmauer, Jamie Maziarz, Mihaela Pavlicev, Ruth Fishman, Lee Koren, Roberto Romero, Günter P. Wagner

## Abstract

Embryo implantation is the first step in the establishment of pregnancy in eutherian (Placental) mammals. Although viviparity evolved prior to the common ancestor of marsupials and eutherian mammals (therian ancestor), implantation is unique to eutherians. The ancestral therian pregnancy likely involved a short phase of attachment between the fetal and maternal tissues followed by parturition rather than implantation, similar to the mode of pregnancy found in marsupials such as the opossum. Embryo implantation in eutherian mammals as well as embryo attachment in opossum, induce a homologous inflammatory response in the uterus. Here, we elucidate the evolutionary mechanism by which the ancestral inflammatory fetal-maternal attachment was transformed into the process of implantation. We performed a comparative transcriptomic and immunohistochemical study of the gravid and non-gravid uteri of two eutherian mammals, armadillo (*Dasypus novemcinctus*) and hyrax (*Procavia capensis*); a marsupial outgroup, opossum (*Monodelphis domestica*); and compared it to previously published data on rabbit (*Oryctolagus cuniculus*). This taxon sampling allows inference of the eutherian ancestral state. Our results show that in the eutherian lineage, the ancestral inflammatory response was domesticated by suppressing a detrimental component *viz*. signaling by the cytokine IL17A, while retaining components that are beneficial to placentation, *viz*. angiogenesis, vascular permeability, remodeling of extracellular matrix. IL17A mediates recruitment of neutrophils to inflamed mucosal tissues, which, if unchecked, can damage the uterus as well as the embryo and lead to expulsion of the fetus. We hypothesized that the uterine decidual stromal cells, which evolved coincidentally with embryo implantation, evolved, in part, to prevent IL17A-mediated neutrophil infiltration. We tested a prediction of this hypothesis *in vitro*, and showed that decidual stromal cells can suppress differentiation of human naïve T cells into IL17A-producing Th17 cells. Together, these results provide a mechanistic understanding of early stages of the evolution of the eutherian mode of pregnancy, and also identify a potentially ancestral function of an evolutionary novelty, the decidual stromal cell-type.

## Introduction

Embryo implantation is the process by which the blastocyst establishes a sustained fetal-maternal interface for the maintenance of pregnancy. It begins with apposition of the blastocyst to endometrial luminal epithelium, followed by its attachment via molecular interactions, and, in many eutherian species, invasion of the endometrium to establish a direct contact with the endometrial connective tissue and vasculature (Mossman 1937, Enders and Schlafke 1969, Schlafke and Enders 1975, Ashary, Tiwari et al. 2018). Implantation is one of the most critical steps in the establishment of a successful pregnancy, but it only occurs in eutherian mammals (also known as Placental mammals). Mammalian viviparity originated before the common ancestor of eutherian mammals and marsupials, i.e. in the stem lineage of therian mammals. However, marsupial and eutherian pregnancies are different in many fundamental ways, including embryo implantation.

Marsupial pregnancy is very short — in most cases shorter than the ovarian cycle (Renfree 1994, McAllan 2011). For most of the duration of marsupial pregnancy, the embryo is present inside of an eggshell (Selwood 2000, Griffith, Chavan et al. 2017) that precludes a direct physical contact between the fetal and maternal tissues. Towards the end of the pregnancy, the eggshell breaks down and the fetal membranes attach to the uterine luminal epithelium. The phase of attachment lasts a short fraction of the length of gestation, and is soon followed by the birth of highly altricial neonates. For instance, pregnancy in the South American marsupial, the grey short-tailed opossum (*Monodelphis domestica*), lasts 14.5 days. Embryo attachment begins approximately on the 12^th^ day post-copulation (dpc) and induces an acute inflammatory response in the uterus (Griffith, Chavan et al. 2017). This inflammation is presumably why embryo attachment precipitates parturition rather than implantation (Chavan, Griffith et al. 2017, Hansen, Faber et al. 2017).

Puzzlingly, in many eutherian mammals such as human, mouse, pig, and sheep, embryo implantation also shows signs of an inflammatory reaction; some of these inflammatory processes are in fact necessary and beneficial for a successful implantation (Keys, King et al. 1986, Barash, Dekel et al. 2003, Waclawik and Ziecik 2007, Plaks, Birnberg et al. 2008, Mor, Cardenas et al. 2011, Dekel, Gnainsky et al. 2014, Robertson and Moldenhauer 2014, Chavan, Griffith et al. 2017, Whyte, Meyer et al. 2017). Resemblance of the physiological process of implantation to an inflammatory reaction appears paradoxical at first because inflammation in later stages of pregnancy leads to the termination of pregnancy. However, analysis of the evolutionary history of embryo implantation suggests that this resemblance is due to the evolutionary roots of implantation in an inflammatory response to embryo attachment (Finn 1986, Griffith, Chavan et al. 2017). Griffith and colleagues (Griffith, Chavan et al. 2017, Griffith, Chavan et al. 2018) argued, based on a comparison of embryo attachment in opossum to implantation in eutherian mammals, that eutherian implantation and the inflammatory marsupial attachment reaction are homologous processes. That is, these processes evolved from an inflammatory fetal-maternal attachment reaction that likely existed in the therian ancestor.

The difference between the fetal-maternal attachment in marsupials and eutherians is its outcome. In the opossum the brief inflammatory attachment results in parturition, whereas in eutherians it results in implantation and establishment of a sustained fetal-maternal interface.

Here, we elucidate the mechanism by which the ancestral attachment-induced inflammatory response was transformed into the process of embryo implantation in the eutherian lineage. We show that the origin of decidual stromal cells (DSC) — a novel eutherian cell type — was integral to this transformation. First, we provide evidence to further support the homology between opossum attachment reaction and eutherian embryo implantation. Then, we compare gene expression in the uterus of opossum during attachment to that in two eutherians during implantation, armadillo and rabbit. The key differences in gene expression suggest that embryo implantation evolved through suppression of a specific module of the ancestral mucosal inflammatory reaction — neutrophil recruitment mediated by the pro-inflammatory cytokine IL17A. We hypothesized that the origin of DSC in the eutherian lineage (Mess and Carter 2006), coincidentally with embryo implantation, was responsible for the suppression of IL17A in members of this clade. A test of this hypothesis using human cells showed that secretions from DSC inhibit the differentiation of Th17 lymphocytes, the primary producers of IL17A, by inducing a non-standard type-1 interferon response that down-regulates their protein synthesis.

## Results and Discussion

### Inflammatory implantation is an ancestral eutherian trait

The inference of homology between the opossum attachment reaction and eutherian embryo implantation is derived from comparison of opossum to species from Boreotheria (Griffith, Chavan et al. 2017). Boreotheria is one of the three major clades that make up Eutheria and includes primates, rodents, ungulates, carnivores, bats and their kin. However, Eutheria also contains two other major clades, Xenarthra and Afrotheria (dos Reis, Inoue et al. 2012, Tarver, dos Reis et al. 2016), for which data on inflammatory gene expression at implantation was previously unavailable. Xenarthra includes armadillo, sloth, anteater, etc; and Afrotheria includes elephant, hyrax, tenrec, aardvark, dugong, etc. See **Figure 1d** for the phylogenetic relationship among eutherian species. To test the inference of homology, we investigated the hypothesis that inflammatory implantation is a shared eutherian character; that is, it is not limited to Boreotheria, but is also observed in Xenarthra and Afrotheria. We did this by assessing the expression of marker genes of inflammation during embryo implantation in the nine-banded armadillo (*Dasypus novemcinctus*) and rock hyrax (*Procavia capensis*), as representatives of Xenarthra and Afrotheria, respectively.

**Figure 1.**
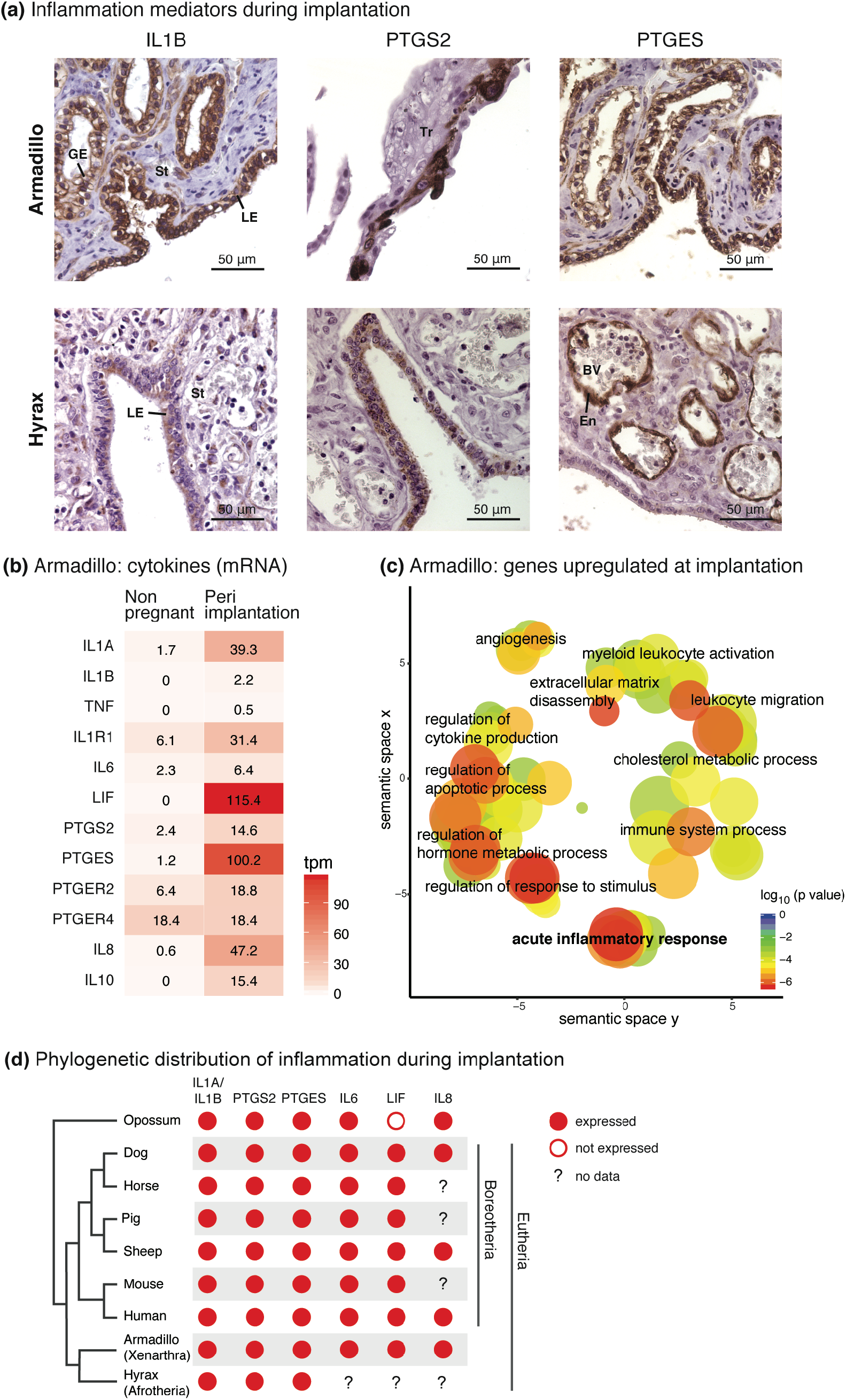
Inflammatory implantation is an ancestral eutherian character. **(a)** Immunohistochemistry for inflammation marker genes IL1B, PTGS2, and PTGES at the fetal-maternal interface in armadillo and hyrax. Nuclei are blue due to hematoxylin counterstaining and the immunostaining signal is brown due to DAB (3,3’-diaminobenzidine). GE = glandular epithelium, LE = luminal epithelium, St = stroma, Tr = trophoblast, BV = blood vessel, En = endothelium. **(b)** Abundance of mRNA transcripts (TPM = Transcripts per Million) of key inflammatory genes in armadillo uterus in non-pregnant and peri-implantation stage. **(c)** Enriched gene ontology (GO) categories among the genes that are upregulated at least 10-fold in armadillo uterus in transition from non-pregnant to peri-implantation stage. GO categories are clustered by semantic similarity. GO categories closer to each other are semantically similar; those represented by red circles are more significantly enriched than those in blue. **(d)** Comparison of expression of inflammatory genes during embryo attachment or implantation in therian mammals. Data for human: cytokines (reviewed in Van Sinderen, Menkhorst et al. 2013), PTGS2 (Marions and Danielsson 1999), PTGES (Milne, Perchick et al. 2001). Data for sheep, horse, pig, dog, and mouse: (reviewed in Chavan, Griffith et al. 2017).

One of the earliest signals of inflammation, IL1B, is expressed during implantation in the luminal epithelium of the endometrium in armadillo and hyrax (**Figure 1a**). PTGS2 and PTGES, enzymes involved in the synthesis of prostaglandin E_2_ (PGE_2_), are also expressed during implantation in both species, although the tissues in which these genes are expressed differ between species. PTGS2 and PTGES are expressed on the two sides of the fetal-maternal interface in armadillo — PTGS2 in the trophoblast, while PTGES in the endometrial luminal epithelium. In hyrax, they are both expressed on the maternal side, with PTGS2 in the luminal epithelium and PTGES in the endothelia within the endometrium.

Next, we used transcriptome data to test whether there is a broad signature of inflammation during implantation in armadillo uterus. A variety of inflammatory genes are up-regulated during armadillo implantation: cytokines such as *IL1A, IL1B, IL6, LIF, IL8* (*CXCL8*), and *IL10*; cytokine receptor *IL1R1*; prostaglandin synthesis enzymes *PTGS2* and *PTGES*; and prostaglandin receptors *PTGER2* and *PTGER4* (**Figure 1b**). None of the cytokines shown in **Figure 1b** is expressed in the non-gravid uterus, i.e. their mRNA abundance is below the operational threshold of 3 TPM (Wagner, Kin et al. 2013), but most of them are expressed highly during implantation. Genes up-regulated more than 10-fold in the transition from non-pregnant to implantation stage are significantly enriched in Gene Ontology (GO) categories related to inflammation (**Figure 1c**), such as acute inflammatory process, immune system process, leukocyte migration, regulation of cytokine production, and regulation of response to stimulus.

We summarized the above expression pattern of inflammation marker genes during implantation on a phylogeny of therian mammals, along with the expression patterns from other representative eutherian species, and from a marsupial, opossum, at the time of attachment between fetal and maternal tissues at 13.5 days post-copulation (dpc) (**Figure 1d**). In all major clades of Eutheria, the uterine changes during embryo implantation closely resemble an acute inflammatory reaction. Parsimoniously, this suggests that embryo attachment-induced uterine inflammatory signaling is a plesiomorphic trait of eutherian mammals, i.e. a trait that was inherited from an ancestral lineage and shared with the sister group, the marsupials, adding further support to the argument that attachment-induced inflammation of marsupials is homologous to eutherian embryo implantation (Griffith, Chavan et al. 2017, Griffith, Chavan et al. 2018, Liu 2018). In other words, eutherian implantation likely evolved from, and through modification of, ancestral attachment-induced inflammation.

### Differences in uterine gene expression between opossum and eutherians

To understand *how* eutherian implantation evolved from the ancestral therian attachment induced inflammation, we compared uterine gene expression during attachment-induced inflammation in opossum (*Monodelphis domestica*) to that during implantation in two eutherian mammals, armadillo (*Dasypus novemcinctus*) and rabbit (*Oryctolagus cuniculus*) (rabbit data from Liu, Zhao et al. 2016). Gene expression patterns shared by armadillo and rabbit are likely to be shared by eutherians in general since armadillo and rabbit are phylogenetically positioned so that their common ancestor is the common ancestor of all extant eutherian mammals.

In the uterine transcriptomes of opossum, armadillo, and rabbit, we classified each gene as either expressed or not expressed (see Methods). We then classified these genes as “opossum-specific” if they are expressed in opossum but not expressed in armadillo and rabbit, and “Eutheria-specific” if they are not expressed in opossum but are expressed in both armadillo and rabbit. There are 446 and 456 genes in these categories, respectively, among the total of 11,089 genes that have one-to-one orthologs in all three species.

First we identified enriched GO categories in the opossum-specific and Eutheria-specific lists of genes (**Figure 2**). To increase the specificity of GO enrichment analysis, we used only the subset of the opossum-specific expressed genes that have an at least 2-fold higher gene expression in the attachment phase compared to the non-pregnant stage (204 genes). Such refinement of the Eutheria-specific gene set was not possible since we do not have a transcriptome of non-pregnant rabbit uterus. The opossum-specific set is enriched for genes related to lipid metabolism, especially mobilization of fatty acids from cell membrane, and lipid transport. Lipid metabolism genes are also upregulated in the pregnant uterus of another marsupial, fat-tailed dunnart (*Sminthopsis crassicaudata*) (Whittington, O’Meally et al. 2018). These genes may have functions related to nutrient transfer or steroid metabolism but lipid metabolism is also important in inflammation: for example, the first step in the synthesis of prostaglandins is to break down membrane triglycerides. The opossum-specific set is also enriched for GO category “cellular response to Interleukin-1”, i.e. genes downstream of IL1A and IL1B. This suggests that although inflammatory signaling is initiated by IL1A and/or IL1B upon embryo attachment in opossum as well as in eutherians, their downstream targets are activated only in the opossum. This observation recapitulates at the molecular level a phenomenon at the organismal level — inflammatory signaling is observed in both, but has different outcomes of parturition and implantation in opossum and eutherians respectively.

**Figure 2.**
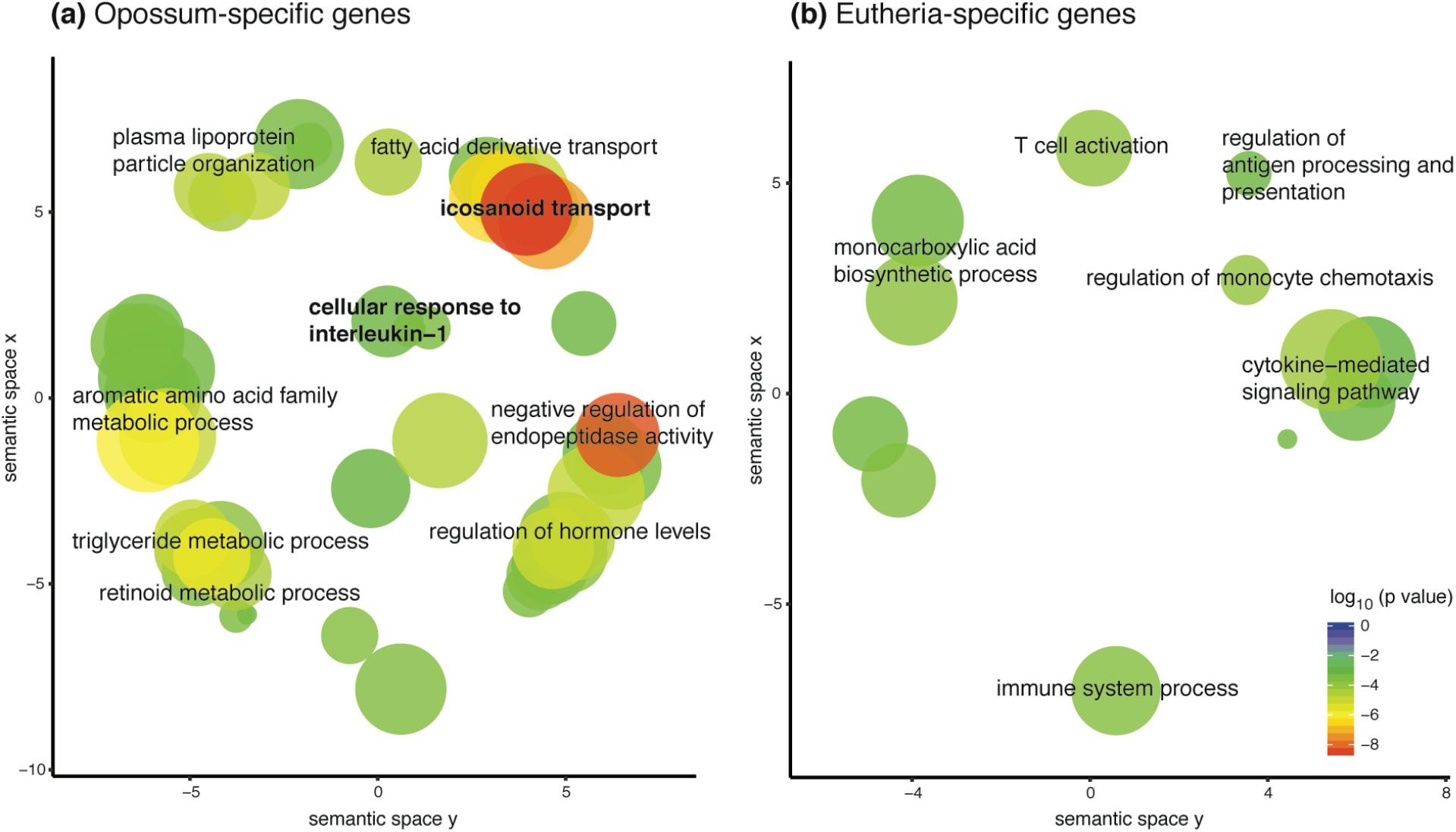
Transcriptomic differences between marsupial and eutherian attachment reaction. Genes were classified as opossum-specific **(a)** and eutherian-specific **(b)**. GO categories enriched in each set are shown in the figures, where semantically similar categories are clustered together in space.

### IL17A expression was suppressed in Eutheria

IL1A and IL1B set off a cascade of inflammatory signaling events mediated by cytokine molecules. Therefore, in order to identify the specific differences between opossum and eutherians in response to interleukin-1, we looked for differences in the expression pattern of cytokines (**Figure 3a**). Cytokines were identified as genes assigned to GO category “cytokine activity” (GO:0005125).

Eutheria-specific expressed cytokines are *CXCL10, CCL5, NDP, WNT7A*, and *LIF*. The first two, CXCL10 and CCL5 attract leukocytes such as T cells, NK cells, and dendritic cells to the sites of inflammation (Schall 1991, Dufour, Dziejman et al. 2002). NDP and WNT7A are both members of the Wnt signaling pathway, which is important for communication between implanting blastocyst and endometrium (Wang and Dey 2006, Chen, Zhang et al. 2009, Sonderegger, Pollheimer et al. 2010); inhibition of this process prevents successful implantation in mouse (Mohamed, Jonnaert et al. 2005). LIF is a critical signaling molecule in eutherian mammals for the differentiation of decidual stromal cells from endometrial stromal fibroblasts; and its expression is therefore obligatory for successful implantation (Shuya, Menkhorst et al. 2011). This set of genes represents cytokines and signaling molecules that were likely recruited within the eutherian lineage to enable decidual cell differentiation, embryo-uterine crosstalk, regulation of leukocyte traffic, and therefore implantation.

**Figure 3.**
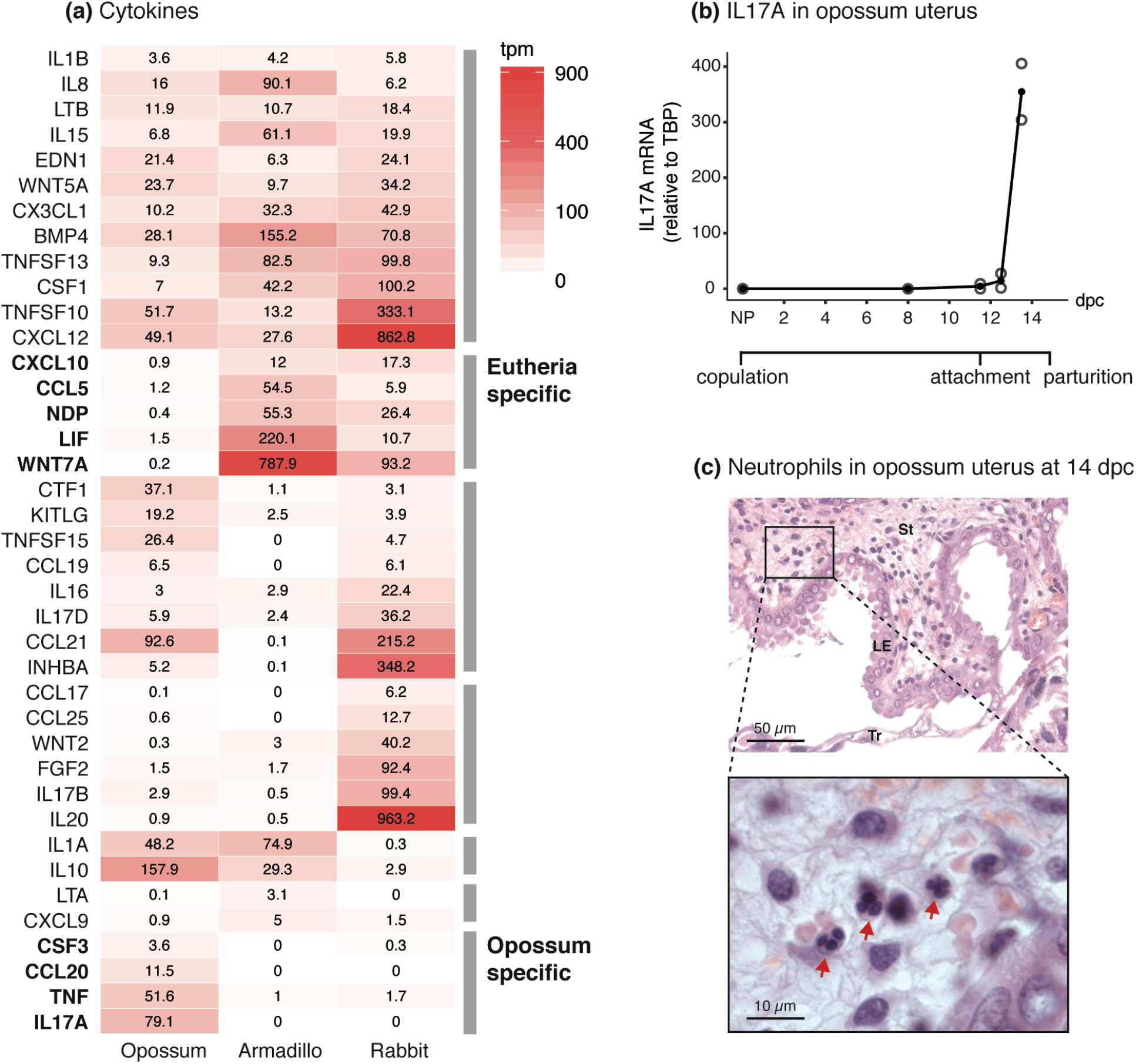
IL17A signaling is suppressed in eutherians. **(a)** Gene expression of cytokines during embryo attachment or implantation in opossum, armadillo, and rabbit uterus. Intensity of red is proportional to the abundance of mRNA. Genes expressed below 3 TPM are considered unexpressed (Wagner, Kin et al. 2013). Rows are ordered by the gene expression patterns across species: expressed in all species, expressed only in Eutheria, expressed in opossum and rabbit, rabbit-specific, expressed in opossum and armadillo, armadillo-specific, and opossum-specific. Cytokines not expressed in any of these species are not shown in the figure. **(b)** Expression of *IL17A* through the pregnancy of opossum, measured relative to *TBP*, by qPCR. Embryo attachment begins around 11.5 days post-copulation (dpc), and pregnancy ends at 14.5 dpc. (number of biological replicates: NP = 3, 8 dpc = 3, 11.5 dpc = 2, 12.5 dpc = 2, 13.5 dpc = 2) (c) Neutrophils infiltration in H&E stained opossum uterus at 14 dpc, indicated by red arrows in the zoomed in micrograph. LE = luminal epithelium, St = stroma, Tr = trophoblast. TPM = Transcripts per Million.

The inverse set, opossum-specific cytokines, includes *CSF3, CCL20, TNF*, and *IL17A*. CSF3, also known as G-CSF (Granulocyte Colony Stimulating Factor) attracts granulocytes (Lieschke, Grail et al. 1994, Panopoulos and Watowich 2008), such as neutrophils, and is positively regulated by IL17A (Ye, Rodriguez et al. 2001, Onishi and Gaffen 2010). CCL20 attracts lymphocytes as well as neutrophils and also is a downstream gene of IL17A (Onishi and Gaffen 2010). This pattern is indicative of a mucosal inflammatory reaction, culminating with recruitment of effector cells such as neutrophils.

Since IL17A is upstream of the other genes in the opossum-specific set, we then measured the expression of *IL17A* in opossum uterus by qPCR to test whether it is induced in response to the attaching embryo or expressed throughout pregnancy. **Figure 3b** shows that IL17A is not expressed in non-pregnant and 8 dpc uteri, and only begins expression at 11.5 dpc, coincidental with the egg-shell breakdown and beginning of fetal-maternal attachment. Its expression reaches a very high level by 13.5 dpc; 79 TPM according to the transcriptomic data. This time-course expression data suggests that *IL17A* is induced specifically in response to embryo attachment in opossum, even though it is completely absent in armadillo and rabbit during implantation (0 TPM in both species). One of the hallmarks of IL17A-mediated inflammation — through the action of cytokines like CXCL8, CSF3, CCL20 — is neutrophil infiltration into the inflamed tissue (Medzhitov 2007, Onishi and Gaffen 2010, Griffin, Newton et al. 2012, Pappu, Rutz et al. 2012, Flannigan, Ngo et al. 2016). Therefore, we tested whether *IL17A* expression in opossum is also accompanied by neutrophil infiltration. Neutrophils are not seen in early stages of pregnancy, but at 14 dpc, neutrophil infiltration is clearly seen histologically (**Figure 3c**). Consistent with the pattern of *IL17A* expression, neutrophils are absent from the fetal-maternal interface at the time of implantation in eutherian mammals (evidence reviewed in Chavan, Griffith et al. 2017).

The absence of *IL17A* expression in eutherian mammals at implantation is likely to be due to its loss in the eutherian lineage rather than its recruitment in the marsupial lineage. The discovery of IL17 homologs as the early-responding cytokines in the sea urchin larva during gut infection (Buckley, Ho et al. 2017) suggests that it may be an ancient mucosal inflammatory cytokine at least as old as deuterostomes. Since the endometrium is a mucosal tissue, IL17A — a key mucosal inflammatory cytokine — is likely to have been expressed in the ancestral therian endometrium during attachment-induced inflammation, and its expression was lost later during evolution in the eutherian lineage.

Because *IL17A* is 1) the most highly expressed gene among opossum-specific cytokines, 2) an important regulator of mucosal inflammation, 3) upstream of chemokines like CSF3, and 4) not expressed in armadillo and rabbit at all, even in a leaky manner, we posited that the loss of *IL17A* expression — and thus the loss of neutrophil infiltration — after embryo attachment was one of the key innovations that transformed the ancestral inflammatory attachment reaction into embryo implantation.

Contrary to the data from armadillo and rabbit, some IL17A expression has been reported at the fetal-maternal interface in mouse and human, specifically in the γδ T cells in the mouse (Pinget, Corpuz et al. 2016). However, functional evidence suggests that even in mouse and human, an induction of IL17A expression at the fetal-maternal interface is detrimental to pregnancy. The strongest evidence perhaps comes from mouse, where injection of polyI:C to mimic viral infection during pregnancy induces the expression of IL17A in the decidua. Maternal IL17A then makes its way into the fetal circulation, interferes with brain development of the embryos, and leads to behavioral abnormalities resembling Autism Spectrum Disorder (ASD) in pups (Choi, Yim et al. 2016). Li and colleagues (Li, Qu et al. 2018) showed that in a mouse model of spontaneous abortion, Th17 cell numbers are higher in the decidua relative to normal pregnancy. In human, Th17 levels in peripheral blood are elevated in women with recurrent spontaneous abortions compared to women with healthy pregnancy (Wang, Hao et al. 2010). Nakashima and colleagues (Nakashima, Ito et al. 2010) found that while IL17A-positive cells were only occasionally present in the deciduae of normal pregnancies, their numbers were significantly elevated in pregnancies with first trimester spontaneous abortions. These studies clearly indicate that IL17A signaling at the fetal-maternal interface is not conducive to successful pregnancy, and the cases in human and mouse where IL17A expression is observed, the downstream signaling is likely inhibited in some way.

Next, we sought to identify the mechanism by which IL17A signaling was suppressed in the evolution of eutherian mammals.

### Decidual stromal cells suppressed IL17A expression at implantation in Eutheria

Decidual stromal cells (DSC) are a novel cell type that originated in the stem lineage of eutherian mammals (Mess and Carter 2006, Wagner, Kin et al. 2014, Erkenbrack, Maziarz et al. 2018). They differentiate from endometrial stromal fibroblasts (ESF) in many eutherian mammals during pregnancy (Gellersen and Brosens 2014) and also during the menstrual cycle in primates, some bats, the elephant shrew (Emera, Romero et al. 2011), and the spiny mouse (Bellofiore, Ellery et al. 2017) in a process called decidualization. The evolution of DSC from ancestral therian ESF was associated with modulation of expression of genes involved in the innate immune response (Kin, Maziarz et al. 2016). DSC perform many functions critical to the maintenance of pregnancy in human and mouse, for example, regulation of the traffic of leukocytes into the endometrium during pregnancy (Nancy, Tagliani et al. 2012, Erlebacher 2013), production of hormones like prolactin, regulation of invasiveness of the trophoblast (Gellersen and Brosens 2014), regulating communication between cell types at the fetal-maternal interface (Pavlicev, Wagner et al. 2017). Outside of euarchontoglirean mammals (primates, rodents and their relatives), however, DSC are not maintained throughout the pregnancy. In bats (Laurasiatheria), hyrax, tenrec (Afrotheria), and armadillo (Xenarthra, also see **Supp. Figure 1**), DSC differentiate around the time of implantation but are often lost soon after implantation. This suggests that the ancestral function of DSC when they originated, was likely to have been limited to the time of implantation (Chavan, Bhullar et al. 2016). Based on their inferred ancestral role at the time of implantation (Chavan, Bhullar et al. 2016), their ability to regulate the immune response during pregnancy (Erlebacher 2013), and because their origin in the eutherian stem lineage (Mess and Carter 2006) coincides with the evolution of suppression of IL17A expression (this study), we hypothesized that DSC played a role in the regulation of IL17A during implantation.

IL17A is typically expressed by Th17 cells, which differentiate from naïve T cells in the presence of IL6 and TGFB1 (Bettelli, Carrier et al. 2006) (**Figure 4a**). We hypothesized that DSC suppress IL17A expression by inhibiting the differentiation of naïve T cells into Th17 cells. To test this hypothesis, we differentiated primary human naïve T cells into Th17, in the presence of DSC-conditioned or control medium and measured IL17A secreted by T cells with ELISA. Treatment of differentiating T cells with DSC conditioned medium decreases their IL17A production significantly (2 fold, p < 10^-6^), while treatment with unconditioned DSC medium has no effect. The decreased level of IL17A is not statistically distinguishable from that in unstimulated naïve T cells, suggesting that DSC-conditioned medium completely suppresses upregulation of IL17A production during Th17 differentiation (**Figure 4b**; see **Supp. Figure 2** for mRNA levels of *IL17A*).

Wu and colleagues (Wu, Jin et al. 2014) showed that Th17 cells are present in the human endometrium during the first trimester and that they are recruited there by DSC, which appears to contradict what we have shown in this study. However, the Th17 cells reported in the decidua by Wu and colleagues are CD45RO+, i.e. they are memory Th17 cells that are already differentiated, while we show that DSC inhibit the differentiation of Th17 cells from naïve T cells.

**Figure 4.**
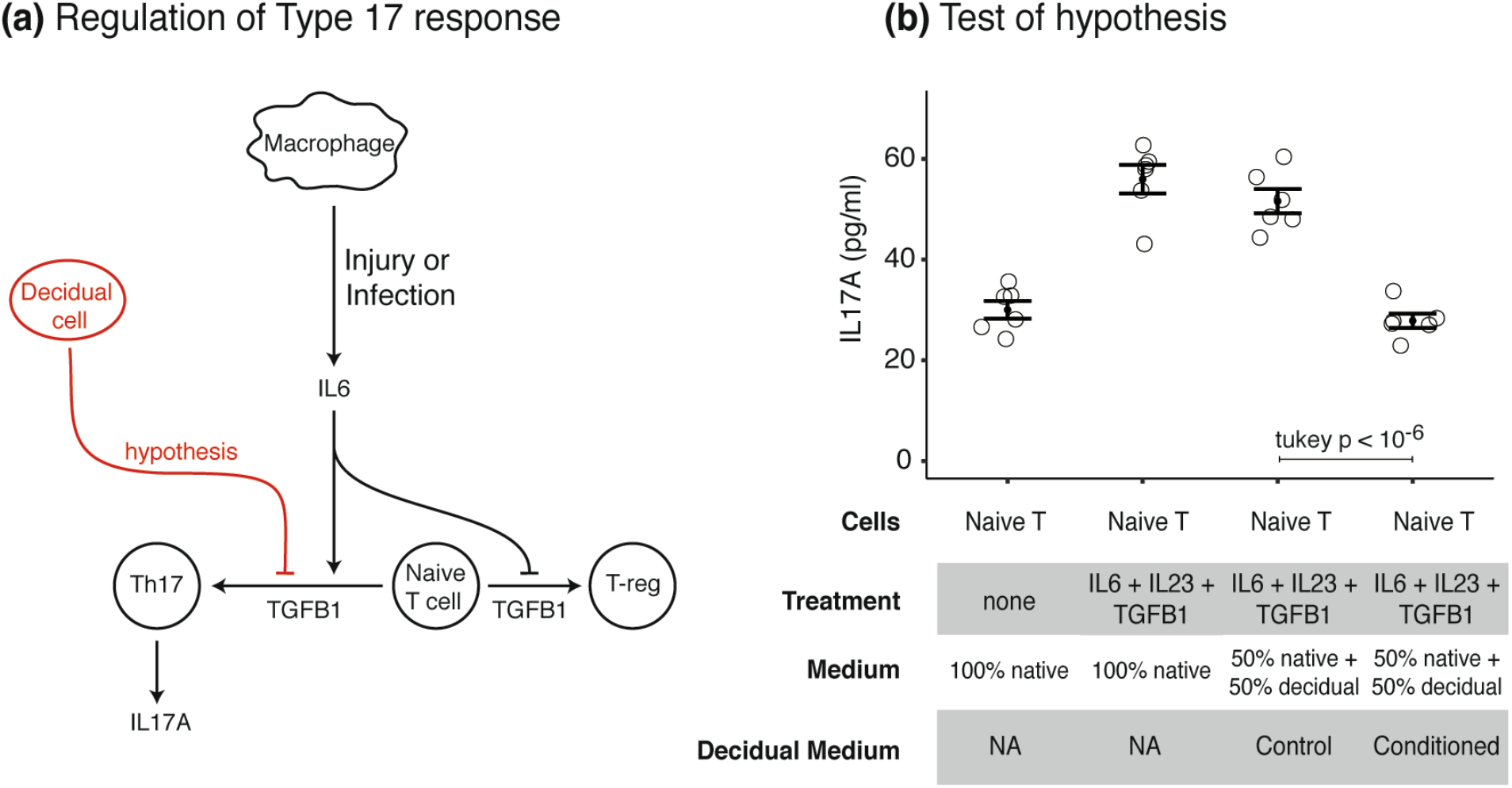
Decidual stromal cells suppress Th17 cell differentiation. **(a)** Schematic showing how Th17 cells differentiate from naïve T cells, and the hypothesis for the role of DSC **(b)** Test of the hypothesis using *in-vitro* differentiation of human naïve T cells into Th17 cells. IL17A secretion (pg/ml) by Th17 cells is shown (mean and standard error of the mean). The first two samples are reference points, where Th17 cells were differentiated in their native medium, and the last two samples were differentiated in DSC control or conditioned medium.

### DSC inhibit protein synthesis during Th17 differentiation

To understand how DSC suppress differentiation of naïve T cells into Th17 cells, we sequenced the transcriptomes of the differentiating T cells that were either treated with DSC conditioned medium or control medium (conditions 3 and 4 from **Figure 4b**). Principal components analysis of the transcriptome data shows that the DSC conditioned medium had a systematic effect on the gene expression profile of the T cells since 60% of the variance between samples can be explained by the first principal component along which the samples separate by treatment (**Figure 5a**).

**Figure 5.**
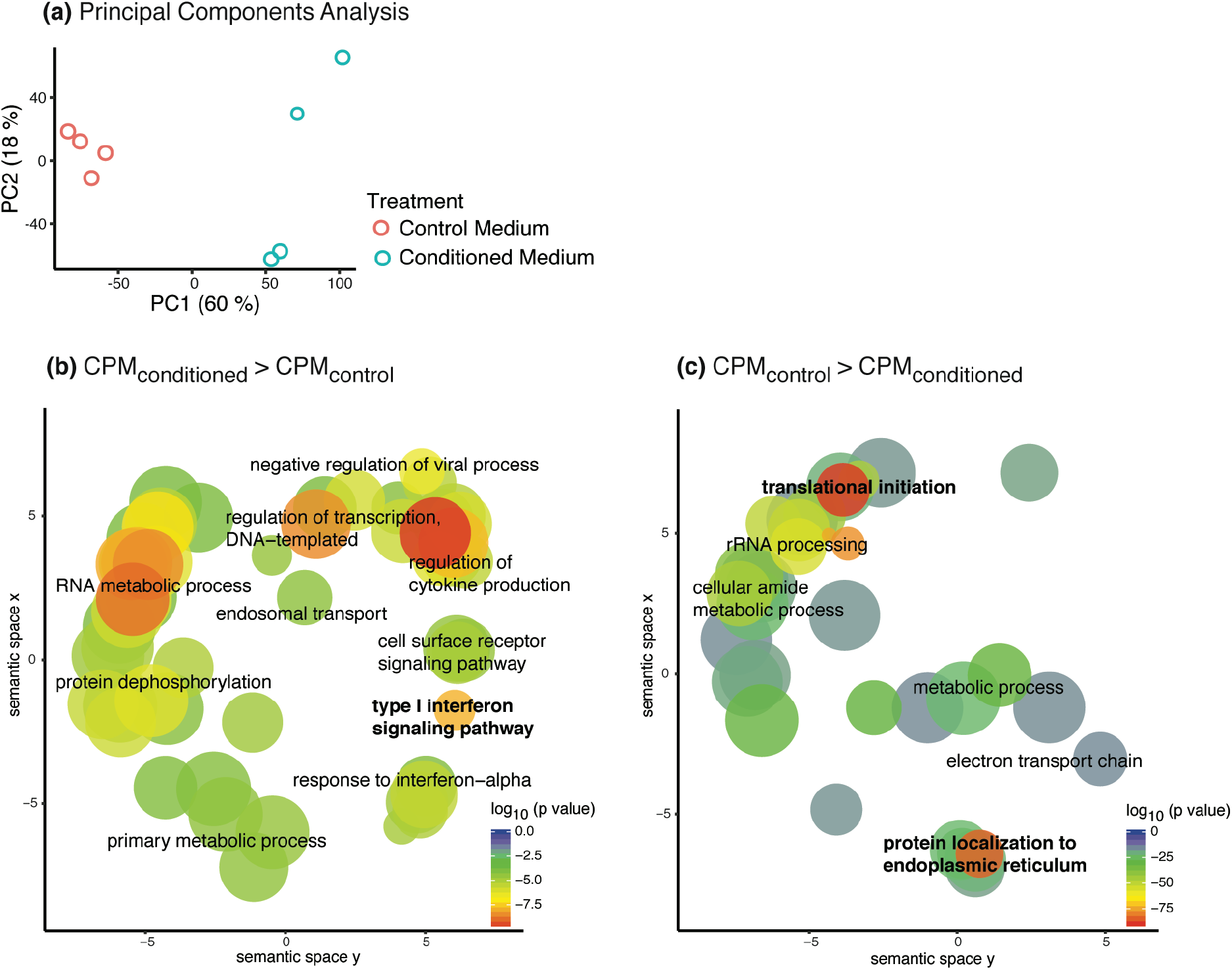
RNA-seq of Th17 cells differentiated in control vs. DSC conditioned medium. **(a)** Principal components analysis of transcriptomes of the control and conditioned medium treated cells. Square root of CPM was used for this analysis. **(b)** and **(c)** respectively show GO categories enriched in genes that have higher expression in T cells treated with conditioned medium compared to the control medium, and vice versa. CPM = Counts per Million.

Genes significantly up-regulated, as assessed by EdgeR (Robinson, McCarthy et al. 2010), in T cells treated with DSC conditioned medium compared to T cells treated with control medium are enriched for GO categories related to defense of viral infection, such as the type-1 interferon signaling pathway, negative regulation of viral process, response to interferon alpha, RNA metabolic process (**Figure 5b**). The down-regulated genes are enriched for GO categories related to protein synthesis and secretion, e.g. rRNA processing, translation initiation, protein localization to endoplasmic reticulum. Genes of the electron transport chain are also down-regulated, suggesting that the cells down-regulated their ATP synthesis and became quiescent (**Figure 5c**).

Then we looked more specifically at genes in the type-1 interferon signaling pathway (**Figure 6a**). Cells infected with virus produce type-1 interferons (IFN-α and IFN-β), which regulate gene expression in an autocrine and paracrine manner via binding to type-1 interferon receptors IFNAR1 and IFNAR2. Curiously, none of the IFN-α genes is expressed in either samples, and IFN-β gene (*IFNB*) is barely expressed in both samples (<1 TPM) but is not differentially expressed in response to DSC conditioned medium. Both IFN receptor genes, however, are significantly up-regulated in samples treated with conditioned medium (**Figure 6b**). Importantly, none of the type-1 interferons are expressed in DSC either (**Figure 6b**). This suggests that the activation of type-1 interferon signaling in T cells is in response to a specific non-interferon signal from DSC and not a genuine viral defense response.

One of the prominent outcomes of type-1 interferon signaling is inhibition of protein synthesis via mRNA degradation and inhibition of translation initiation machinery (Kindt, Goldsby et al. 2007). This ensures that viral nucleic acids do not replicate within the host cells and produce more viral particles. In T cells treated with DSC-conditioned medium, three out of four 2’-5’ oligoadenylate synthase genes (*OAS1, OAS2* and *OAS3*) genes are up-regulated (**Figure 6c**). These genes activate RNaseL, which in turn degrades mRNA. Most of the eukaryotic translation initiation factors (EIFs) are also down-regulated and at the same time two negative regulators of EIFs, *viz*. *EIF2AK2* and *EIF4EBP2* are up-regulated (**Figure 6d**), indicating decreased protein synthesis.

Together these observations suggest that DSC suppress Th17 differentiation by inducing type-1 interferon signaling, which consequently decreases overall protein synthesis and ATP synthesis. However, DSC secrete neither type-1 IFNs nor do they induce the expression of type-1 IFNs in T cells, suggesting that DSC hijack the viral defense mechanism downstream of the IFN receptors to inhibit protein synthesis in T cells. This is in contrast to the bovine pregnancy in which the ruminant-specific type-1 interferon IFN-τ (*IFNT*) from the embryo suppresses *IL17A* expression in maternal peripheral blood mononuclear cells (Talukder, Rashid et al. 2018). Signals from DSC that inhibit protein synthesis in human T cells are unlikely to be generic molecules that can induce type-1 IFN signaling, e.g. cell free RNA or DNA. First of all, such molecules would induce type-1 IFN signaling in all cells at the fetal-maternal interface, which would be detrimental to the maintenance of pregnancy, just as a viral infection is (Yockey and Iwasaki 2018, Yockey, Jurado et al. 2018). A generic signal would also suppress differentiation of naïve T cells into not only Th17 cells but also other effector T cells. However, T-reg cells are necessary for maintenance of pregnancy (Aluvihare, Kallikourdis et al. 2004), and their differentiation should not be hindered by DSC. Potential signals from DSC that may be acting on T cells are discussed in the supplementary material (**Supp. Figure 3**).

**Figure 6.**
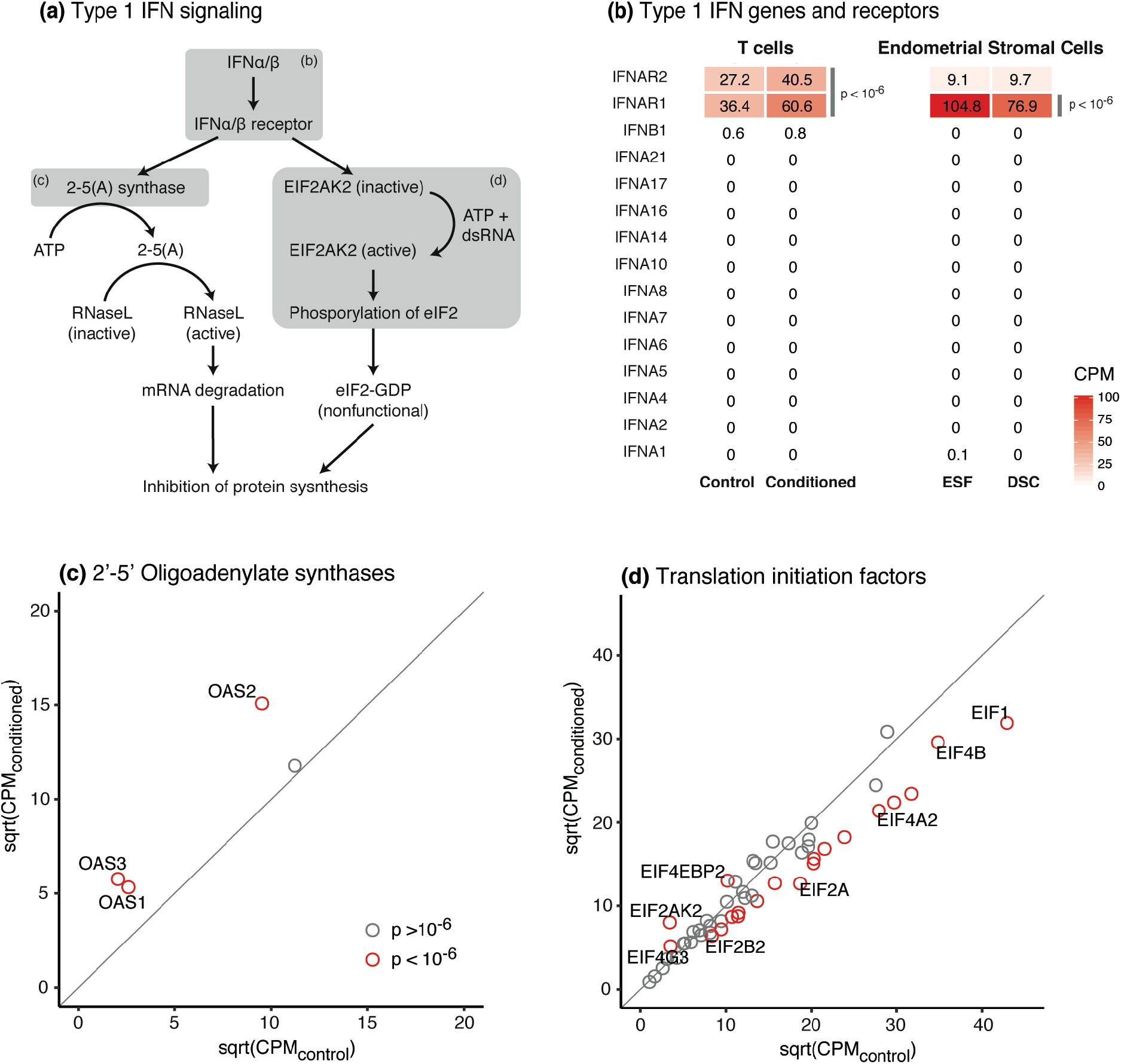
Interferon signaling genes in T cells treated with DSC conditioned medium. **(a)** Schematic showing how type-1 interferon signaling pathway inhibits protein synthesis, adapted from (Kindt, Goldsby et al. 2007). Grey boxes show the sets of genes from this pathway whose gene expression is shown in the following panels of the figure. **(b)** Type-1 interferon genes and their receptors in Th17 cells differentiated with control or DSC conditioned medium, and in endometrial stromal cells. **(c)** 2’-5’ Oligoadenylate synthase genes. **(d)** Eukaryotic translation initiation factors and some of their regulators. Red points in (c) and (d) represent genes significantly differentially expressed between control and DSC conditioned medium treated T cells according to EdgeR (see Methods). CPM = Counts per Million.

### A model for the evolution of implantation

Placing the results from this study in the context of previous studies, the following model for the evolution of embryo implantation emerges (**Figure 7**).

**Figure 7.**
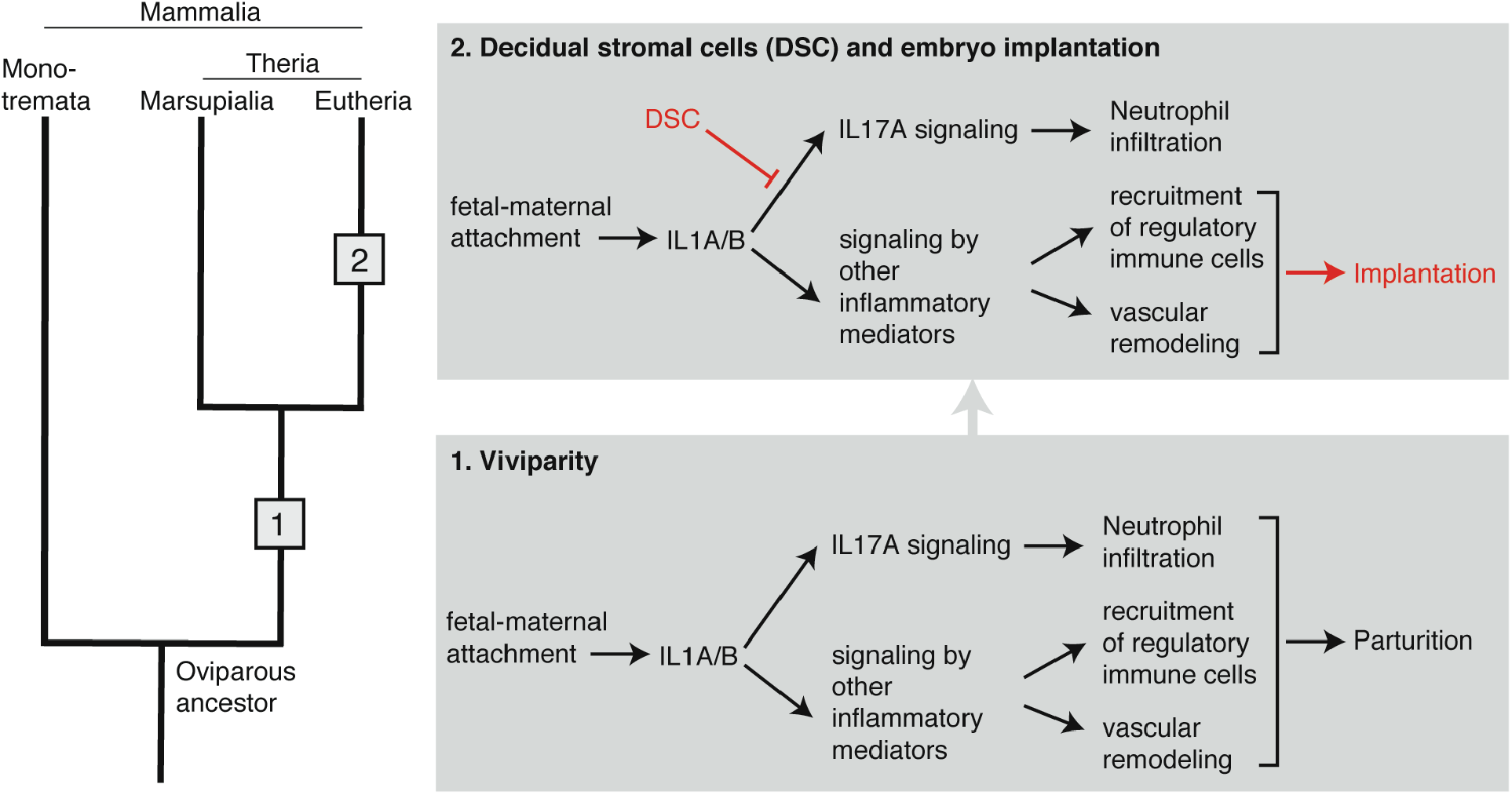
Model for the evolution of eutherian implantation from attachment-induced inflammation. Evolutionary events in the therian and eutherian stem lineages are represented by numbers 1 and 2 respectively.

The ancestor of all mammals was an egg-laying amniote. Mammalian viviparity originated in the stem lineage of therian mammals by early “hatching” of the embryo while it was still within the uterus/shell gland, leading to a direct physical contact between fetal membranes and the uterine endometrial lining. This novel tissue interaction, and potentially an irritation of the endometrial lining from fetal proteases that help dissolve the shell (Griffith, Chavan et al. 2017), induced an acute inflammatory response in the endometrium. An inflamed uterus, unable to maintain and nourish a live embryo within it, expelled the embryo, i.e. parturition ensued as a consequence of attachment-induced inflammation.

Hence we assume that the ancestral condition for therians was a typical mucosal inflammation in response to embryo attachment. However, in the stem lineage of eutherian mammals, this inflammatory response became evolutionarily modified such that the endometrium could maintain and nourish the embryo, even though it initiated an inflammation-like response. One of these modifications was the suppression of IL17A signaling, which resulted in the prevention of recruitment of neutrophils to the endometrium during implantation. This may have been a crucial modification to the inflammatory response since neutrophils are known to cause collateral tissue damage from the digestive enzymes they secrete; and the attaching embryo also would likely have not been spared from this tissue damage. Turning off IL17A early in the evolution of the eutherian lineage may have allowed the endometrium to maintain the embryo for a prolonged period. In contrast, many other aspects of the ancestral inflammation may have been beneficial to the maintenance and nourishment of the embryo, e.g. increased vascular permeability and angiogenesis may have helped nutrient transfer, therefore still maintained as a necessary part of implantation. The ancestral attachment-induced inflammation induced a stress reaction in the endometrial stromal fibroblasts (ESF), causing the death of most of these cells. In the eutherian lineage, however, ESF evaded the stress-induced cell death (Kajihara, Jones et al. 2006, Leitao, Jones et al. 2010, Muter and Brosens 2018) by evolving mechanisms to differentiate into a novel cell type, decidual stromal cells (DSC) (Erkenbrack, Maziarz et al. 2018). It is this novel cell type that likely brought about the suppression of IL17A by inhibiting the differentiation of IL17A-producing cells at the fetal-maternal interface, and thus enabled the evolution of embryo implantation and a sustainable fetal-maternal interface.

## Methods

### Animals and tissue samples

Nine-banded armadillos (*Dasypus novemcinctus*) were collected in Centerville, TX, USA, in accordance with Yale University IACUC protocol #2014–10906. Two females were used in this study — one non-pregnant and one at the peri-implantation stage of pregnancy where the fetal membranes have begun invading the endometrium but no placental villi are yet formed. Rock hyrax (*Procavia capensis*) samples were collected at Bar-Ilan University, Israel. One out of three females collected was in the implantation phase of pregnancy as determined by histological examination. The blastocyst was attached to the uterine lumen but had not started invading the endometrium. Opossum (*Monodelphis domestica*) tissues were collected from the colony housed at Yale University. Samples were fixed in 4% paraformaldehyde in phosphate-buffered saline (PBS) for histology and immunohistochemistry, saved in RNAlater (Ambion) for RNA extraction. For more information about armadillo samples, see (Chavan and Wagner 2016), and about opossum samples see (Griffith, Chavan et al. 2017). A list of animals used in this study is given in **Supp. Table 1**.

### Immunohistochemistry

Formaldehyde-fixed tissues were dehydrated in ethanol, cleared in toluene, and embedded in paraffin. Sections of 5µm thickness were made on a microtome and placed on poly-L-lysine coated glass slides. Immunohistochemistry was performed by following the protocol in (Chavan and Wagner 2016). Briefly, slides were incubated at 60 °C for 30 min and allowed to cool at room temperature for 5 min. Paraffin was removed by de-waxing the slides in xylene. Slides were then rehydrated by successive washes in 100% ethanol, followed by running tap water. Sodium Citrate buffer (pH 6.0) was used for heat mediated antigen retrieval. After washing the slides in PBS, they were blocked in a 0.1% solution of Bovine Serun Albumin (BSA) in PBS.

Endogenous peroxidases were suppressed with Peroxidase Block (DAKO). Optimized dilutions of primary antibodies (see **Supp. Table 2**) were added to the slides and were incubated overnight at 4°C in a humidification chamber. Secondary antibody was added after washing the primary antibody off with PBS and 0.1% BSA-PBS, and incubated for 1 hour at room temperature, and washed off with PBS and 0.1% BSA-PBS. Horseradish peroxidase (HRP) tagged secondary antibodies were detected by either 3,3’-diaminobenzidine (DAB) and counterstained with hematoxylin.

### Cell culture experiments

#### DSC and DSC conditioned medium

Immortalized human endometrial stromal fibroblasts (hESF) from ATCC (ATCC; cat. no. CRL-4003) were cultured in growth medium with the following contents per liter: 15.56 g DMEM (D2906, Sigma-Aldrich), 1.2 g sodium bicarbonate, 10 ml sodium pyruvate (11360, Thermo Fisher), 10 ml ABAM (15240062, Gibco), 1 ml ITS (354350, VWR), and 100 ml charcoal-stripped FBS. ESF were differentiated into decidual stromal cells (DSC) in differentiation medium with the following contents per liter: 15.56g DMEM (D8900, Sigma-Aldrich), 1.2g sodium bicarbonate, 10 ml ABAM, 0.5 mM cAMP analog 8-Bromoadenosine 3′,5′-cyclic monophosphate sodium salt (B7880, Sigma-Aldrich), 1 µM progesterone analog Medroxyprogesterone 17-acetate (MPA; M1629, Sigma-Aldrich), and 20 ml FBS.

Cells were differentiated for 48 hours at 37 °C, conditioned medium was collected from at least 6 replicate flasks, filtered through sterile 0.45 µm filter to remove cell debris, aliquoted, and frozen at −80 °C until used. For control samples, decidualization medium was incubated for 48 hours at 37 °C without any cells in at least 6 replicate flasks, processed in the same way as conditioned medium, and frozen at −80 °C.

#### T cells

Human Peripheral Blood Mononuclear Cells (PBMC) were isolated from whole blood (70501, StemCell Technologies) using Lymphoprep (07851, StemCell Technologies) and 50 ml SepMate tubes (85450, StemCell Technologies). These PBMCs were used to isolate CD4+ CD45RO− naïve T cells with EasySep Human Naïve CD4+ T Cell Isolation Kit (19555, StemCell Technologies). Naïve T cells were resuspended at 10^6^ cells/ml in ImmunoCult-XF T Cell Expansion medium (10981, StemCell Technologies) in the presence of ImmunoCult CD3/CD28/CD2 T Cell Activator (10970, StemCell Technologies) for culture or Th17 differentiation. For Th17 differentiation the following were added to the above cell suspension: 20 ng/ml IL6 (78050, StemCell Technologies), 5 ng/ml TGFB1 (78067, StemCell Technologies), and 50 ng/ml IL23 (14–8239-63, eBioScience). If conditioned medium was used, the above suspension was made in 1:1 solution of conditioned or control medium and the T cell expansion medium. Cell suspension was then transferred to 24-well plates, with 10^6^ cells per well. Samples with different treatments were placed in a Latin square design on 24-well plates to prevent systematic effects arising from the positions of the samples in the plate. Cells were incubated at 37 °C for 7 days for differentiation. Cells, which are in suspension, were spun down to collect the supernatant and cell pellet. RNA was extracted from the cell pellets for RNA-seq (see below), and secreted IL17A was measured in the supernatants with Quantikine ELISA kit for human IL17A (D1700, R&D Systems).

### RNA-seq data

Among the animal tissue collected for this study, armadillo samples were used for RNA-sequencing. RNA was extracted from whole uteri. Sequencing library for RNA from non-pregnant armadillo was prepared and sequenced at the Yale Center for Genome Analysis. Library preparation and sequencing of the RNA from implantation stage armadillo was performed at Cincinnati Children’s Hospital Medical Center.

RNA from CD4+ naïve T cells differentiated into Th17 cells in the presence of DSC conditioned medium was extracted using Qiagen RNeasy Micro Kit (74004, Qiagen). Libraries were prepared and sequenced at the Yale Center for Genome Analysis.

Data for rabbit uterus (*Oryctolagus cuniculus*), grey short-tailed opossum (*Monodelphis domestica*) and human endometrial stromal cells were downloaded from Gene Expression Omnibus (GEO) (Barrett, Wilhite et al. 2013). GEO Accession numbers of the downloaded datasets as well as those generated in this study are listed in **Supp. Table 3**.

### RNA-seq analysis

RNA-seq data were aligned to the following Ensembl genomes using TopHat2 (Kim, Pertea et al. 2013): GRCh37 for human, DasNov3 for armadillo, OryCun2 for rabbit, and MonDom5 for opossum. Number of reads mapping to genes were counted with HTSeq (Anders, Pyl et al. 2015). Read counts were normalized to Transcripts per Million (TPM) (Wagner, Kin et al. 2012), and 3 TPM was used as an operational threshold to call genes as expressed or unexpressed (Wagner, Kin et al. 2013). Median length of all transcripts of a gene was used as its ‘feature length’ for TPM normalization. Differential gene expression analysis for human T cell and endometrial stromal cell data was performed on protein coding genes with the R package EdgeR (Robinson, McCarthy et al. 2010). For these samples, read counts are normalized by EdgeR to Counts per Million (CPM), and are presented as such.

In analyses involving multiple species, only those genes were included that have one-to-one orthology among the species compared. Orthology data from Ensembl Compara database (Herrero, Muffato et al. 2016) were used. In these analyses, read counts were re-normalized to TPM using only the set of orthologous genes in order to make TPM values comparable between species.

Enriched gene ontology (GO) categories in sets of genes were identified with the online tool GOrilla (Eden, Navon et al. 2009). Lists of enriched GO categories were visualized on REViGO (Supek, Bošnjak et al. 2011), which clusters semantically similar GO terms in space, simplifying interpretation of long lists of redundant GO categories.

### qPCR

Expression of IL17A through opossum pregnancy was measured using qPCR at a finer time scale than the trascriptomic data. RNA from non-pregnant, 8 dpc, 11.5 dpc, 12.5 dpc, and 13.5 dpc uteri was extracted and reverse transcribed to cDNA using High Capacity Reverse Transcriptase Kit (4368814, Thermo Fisher). Abundance of IL17A mRNA was measured relative to TBP (Tata Binding Protein) mRNA with a standard curve approach using Power SYBR Green PCR Master Mix (4367659, Applied Biosystems) on StepOne Plus Real Time PCR System (Applied Biosystems). Primer sequences are in **Supp. Table 4**.

## Acknowledgements

This research was supported by John Templeton Foundation Grant #54860 to GPW. We would like to thank Haleigh Larson for assistance with histology.

